# TMS provokes target-dependent intracranial rhythms across human cortical and subcortical sites

**DOI:** 10.1101/2023.08.09.552524

**Authors:** Ethan A. Solomon, Jeffrey B. Wang, Hiroyuki Oya, Matthew A. Howard, Nicholas T. Trapp, Brandt D. Uitermarkt, Aaron D. Boes, Corey J. Keller

## Abstract

Transcranial magnetic stimulation (TMS) is increasingly deployed in the treatment of neuropsychiatric illness, under the presumption that stimulation of specific cortical targets can alter ongoing neural activity and cause circuit-level changes in brain function. While the electrophysiological effects of TMS have been extensively studied with scalp electroencephalography (EEG), this approach is most useful for evaluating low-frequency neural activity at the cortical surface. As such, little is known about how TMS perturbs rhythmic activity among deeper structures – such as the hippocampus and amygdala – and whether stimulation can alter higher-frequency oscillations. Recent work has established that TMS can be safely used in patients with intracranial electrodes (iEEG), allowing for direct neural recordings at sufficient spatiotemporal resolution to examine localized oscillatory responses across the frequency spectrum. To that end, we recruited 17 neurosurgical patients with indwelling electrodes and recorded neural activity while patients underwent repeated trials of single-pulse TMS at several cortical sites. Stimulation to the dorsolateral prefrontal cortex (DLPFC) drove widespread low-frequency increases (3-8Hz) in frontolimbic cortices, as well as high-frequency decreases (30-110Hz) in frontotemporal areas, including the hippocampus. Stimulation to parietal cortex specifically provoked low-frequency responses in the medial temporal lobe. While most low-frequency activity was consistent with brief evoked responses, anterior frontal regions exhibited induced theta oscillations following DLPFC stimulation. Taken together, we established that non-invasive stimulation can (1) provoke a mixture of low-frequency evoked power and induced theta oscillations and (2) suppress high-frequency activity in deeper brain structures not directly accessed by stimulation itself.

## Introduction

Transcranial magnetic stimulation (TMS) has been heralded as a transformative technology in the treatment of neuropsychiatric illness. Through the induction of a magnetic field via a coil placed near the skull, TMS allows physicians to target specific brain regions and networks for modulation, completely non-invasively and often in outpatient clinical settings. Of particular interest has been its application in psychiatry, where it is effective for depression and shows promise for bipolar, addiction, and other psychiatric disorders. It was recently found that TMS delivered to individually-selected brain areas multiple times per day can yield rapid and sustained improvement in depressive symptoms (*1*, *2*).

These clinical successes suggest that repetitive stimulation of cortical targets may change neural activity at the circuit- or network-level of brain function. Invasive measures in animal models have shown TMS can induce neural spiking and alter properties of the local field potential, among other modulatory effects (*3*, *4*). In humans, investigators have long probed motor-evoked potentials (*5*), and more recently used fMRI BOLD (*6*) and scalp EEG (*7*, *8*) to characterize changes in TMS-induced neural activity and plasticity. In general, such studies found that repetitive TMS can alter cortical excitability by variably promoting long-term potentiation (LTP) or depression (LTD)-like effects – depending on the stimulation parameters(*5*, *9*) – as well as drive changes in neural activity in distant brain regions through functional or structural connections (*10*–*13*). A hypothesized antidepressant effect of TMS, for example, arises from strengthened connectivity between the dorsolateral prefrontal cortex (DLPFC) and downstream regions within a cognitive-control network (*2*, *14*). While foundational to our understanding of TMS neurophysiology, these non-invasive studies are fundamentally limited in their power to describe brain dynamics on the detailed spatiotemporal scale necessary to understand the neural effects of TMS and guide novel treatments.

In parallel, research in neurosurgical patients investigates neural dynamics on a finer scale, leveraging the ability to electrically stimulate and record from intracranial electrodes (intracranial EEG; iEEG). Such indwelling electrodes record neural signals with precise spatial localization, high temporal resolution, low noise, and can access deep brain structures which are difficult to measure through scalp EEG (*15*). In these studies, direct cortical electrical stimulation (DES) has been shown to provoke widespread, rhythmic changes in neural activity that appear similar to the spontaneous, endogenous oscillations which are crucial to sensory and cognitive processing (*16*–*20*). Moreover, stimulation-provoked oscillations are observed in deeper structures – like the hippocampus and amygdala – that play important roles in cognition and disease pathogenesis (*19*, *21*).

Although neural oscillations are critical in neuropsychiatric disease, we have only a limited understanding of how TMS affects such oscillatory neural activity (*19*, *20*). Studies combining scalp EEG recordings and TMS tend to focus on the evoked responses (termed TMS-evoked potentials, or TEPs (*10*, *22*–*27*)), which are phase-locked to stimulation and are typically thought of as a signature of bottom-up propagation of activity. Some investigators have attempted to differentiate these evoked responses from “true” oscillations that occur with variable phase relation to the stimulus (*28*–*31*) (also called “induced”), as these may reflect higher-order cognitive processing. However, making this distinction can be theoretically and methodologically ambiguous (*32*, *33*), and there is no clear consensus on how TMS affects these two neuronal processes. Additionally, scalp EEG offers unreliable estimates of oscillatory activity at higher frequencies due to signal attenuation by the skull, muscle artifact, and the generally smaller spatial extent of higher-frequency signals(*34*). Finally, scalp studies are limited in their ability to anatomically localize the origin of a neural event, particularly for areas far from the cortical surface (*10*, *26*, *27*, *35*, *36*).

Until recently, TMS in neurosurgical patients with indwelling electrodes was avoided for safety concerns, though new evidence suggests the approach is safe. Specifically, Wang and colleagues demonstrated in phantom brains that TMS does not displace, induce thermal changes, or elicit substantial changes in electrical fields in implanted electrodes. Moreover, in neurosurgical patients they observed intracranial TMS-evoked potentials (iTEPs) in functionally connected regions that were downstream from a cortical target, demonstrating invasively-recorded neural responses to non-invasive stimulation (*37*, *38*). While an important advance, this study did not evaluate the rich time-frequency information that can be extracted from intracranial electrodes with both high fidelity and spatiotemporal resolution. Such precise time-frequency information would allow investigators to tease out neural effects at specific frequencies of interest, a key physiological dimension which may be foundational to the brain’s functioning (*39*, *40*).

To that end, we set out to characterize the spectral response of the human brain to TMS. We recruited 17 neurosurgical patients with indwelling cortical grids, strips, and depth electrodes while patients underwent repeated single-pulse trials of TMS. By comparing to sham stimulation, we quantified TMS-related spectral responses, particularly focusing on the theta (3-8Hz) and gamma (30-50Hz) bands that have been implicated in cognition and neuropsychiatric disease (*41*), as well as the high-frequency activity (HFA; 70-110Hz) range that may reflect population spiking (*42*). Our goals were to build on existing TMS-EEG and TMS-fMRI work by (1) understanding whether TMS can induce neural oscillations, as opposed to evoked responses, (2) examining the high-frequency spectral responses to TMS, and (3) characterizing the responses of subcortical structures to cortical stimulation. We hypothesized that the effects of TMS would be most prominent in cortical areas local to stimulation itself, but that signals would also propagate to deeper structures and be detectable principally as a burst of broadband spectral power consistent with an evoked potential.

We found that TMS was generally associated with widespread increases in low frequency power that was strongest within 500ms of each pulse. DLPFC stimulation drove increases in low-frequency power in frontolimbic cortices, but also yielded a longer-lasting suppression of gamma and HFA power in the lateral and medial temporal lobes. Parietal stimulation provoked low-frequency power increases specifically in the medial temporal lobe (MTL). Phase analysis demonstrated elevated phase locking of low-frequency power following stimulation – consistent with an evoked response – but a trial-level analysis revealed induced theta oscillations within anterior frontal regions, specifically following DLPFC stimulation. Taken together, these results demonstrate that TMS can be used to modulate the neural activity of cortical and subcortical structures, provoking a mixture of evoked responses and induced theta oscillations, while also suppressing higher-frequency content lasting several hundred milliseconds following each pulse.

## Results

Combined TMS and iEEG allows for a detailed spectrotemporal analysis of TMS-related neurophysiology that is inaccessible to non-invasive measurements. Briefly, we recruited 17 neurosurgical subjects with indwelling electrodes who underwent single-pulse active TMS (spTMS; *n*=50-150 trials, 0.5Hz) and sham (*n*=50-300 trials), targeting the DLPFC or parietal cortex (**Figure 1A**). Target sites were selected using MRI-guided neuronavigation; DLPFC was directed to the Beam F3 target, while parietal sites were selected by the anatomical subregion with maximal resting state fMRI-based connectivity to the hippocampus (*43*) (see *Methods* for details). By statistically comparing the spectral power following sham and TMS pulses, we mapped the frequency-domain responses to stimulation across a wide array of intracranially-recorded brain regions, including bilateral frontal, temporal, parietal, limbic, and medial temporal areas (**Figure 1B, 1C**).

**Figure 1.**
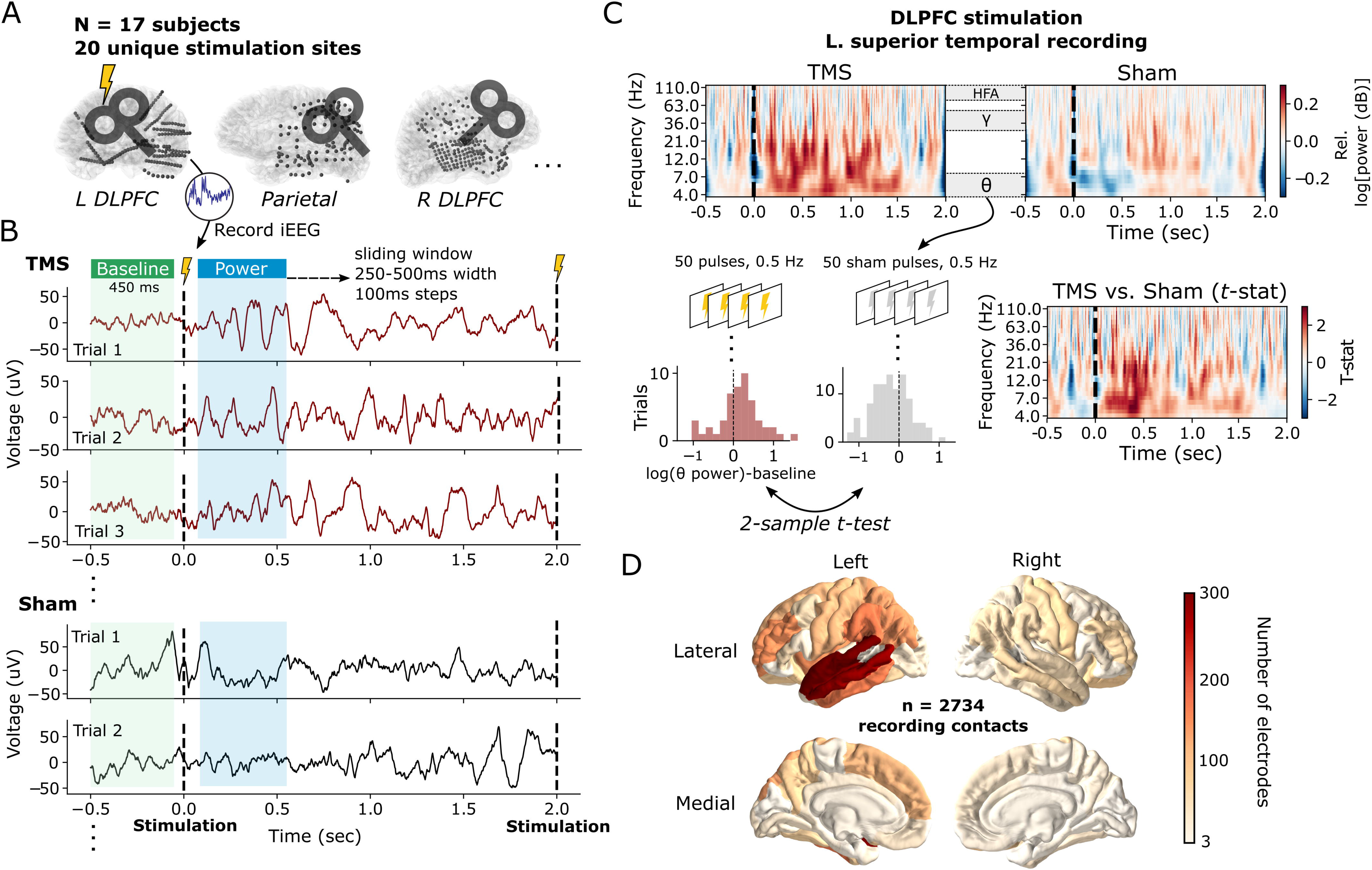
Stimulation protocol and analysis pipeline in an example subject. **(A)** Example schematic representations of the intracranial recording locations and stimulation locations in three subjects from the 17-subject dataset. **(B)** Data shown for one superior temporal gyrus (STG) stereo-EEG recording site, in one subject, recorded during TMS stimulation of the left DLPFC. For each stimulation site within a subject, single-pulse TMS was delivered at least 50 times with 2-second inter-stimulation intervals. In a separate block, at least 50 sham pulses were delivered by flipping the direction of the coil away from the skull, eliminating the induced electric field within the brain, while keeping other experimental parameters constant. For all recording contacts, intracranial EEG was simultaneously recorded and multitaper spectral power extracted in 500ms intervals (250ms for HFA to capture faster fluctuations) starting 50ms after the stimulation pulse (see *Methods* for details). To correct for changes in baseline power from trial-to-trial, power was also computed preceding each stimulation event, and subtracted from post-stimulation power. **(C)** Example time-frequency spectrograms of the power response to TMS and sham stimulation (*top row*), derived from the same superior temporal gyrus iEEG data as in (B). Throughout this study, the power measured across TMS trials was statistically compared to the power following sham trials using a 2-sample *t-*test, generating a *t-*statistic which reflects the degree to which TMS increases or decreases oscillatory power relative to sham (*bottom row*). Sample data is shown for power extracted from the theta (3-8Hz) band within the first 500ms following stimulation. Full spectrograms are shown for completeness; statistical comparisons are rendered on data that excludes the stimulation pulse itself as described in (B). **(D)** Total count of recording contacts (n = 2374) for each Desikan-Killiany-Tourville (DKT) cortical parcellation in the 17-subject dataset.

### Brain-wide spectral responses to DLPFC and parietal stimulation

We first took an overarching view of the brain’s response to TMS, asking if significant differences between TMS and sham-related spectral power were observable in the theta, gamma, or HFA bands across lobe-level regions-of-interest (**Supplemental Figure 1, Figure 2A**). By measuring spectral power in successive windows following stimulation (see *Methods*), we used linear mixed-effects modeling (LMM) to find significant early (starting 50ms post-stimulation in 500ms windows) increases in theta power from DLPFC stimulation, specifically in the frontal (Wald test, *z* = 3.20, *P* = 0.001, Intercept = [0.283, 1.179] 95% CI) and limbic cortices (Wald *z* = 4.161, *P* < 0.001, Intercept = [0.327, 0.910] 95% CI). This effect became nonsignificant in time windows starting 150ms after stimulation in both regions. A significant early theta response (50-550ms) was also observed in the MTL following parietal stimulation (Wald *z* = 3.75, *P* < 0.001, Intercept = [0.199, 0.636] 95% CI). DLPFC stimulation caused an anatomically broad suppression of activity in the gamma and HFA bands (starting 250-450ms post-stimulation), which only survived FDR correction in the temporal cortex (Wald *z* = -3.81, *P* < 0.001, Intercept = [-0.50, -0.16] 95% CI). Of note, by using a windowed spectral approach beginning 50ms after stimulation, we avoid the possibility of direct contamination by the stimulation artifact itself.

**Figure 2.**
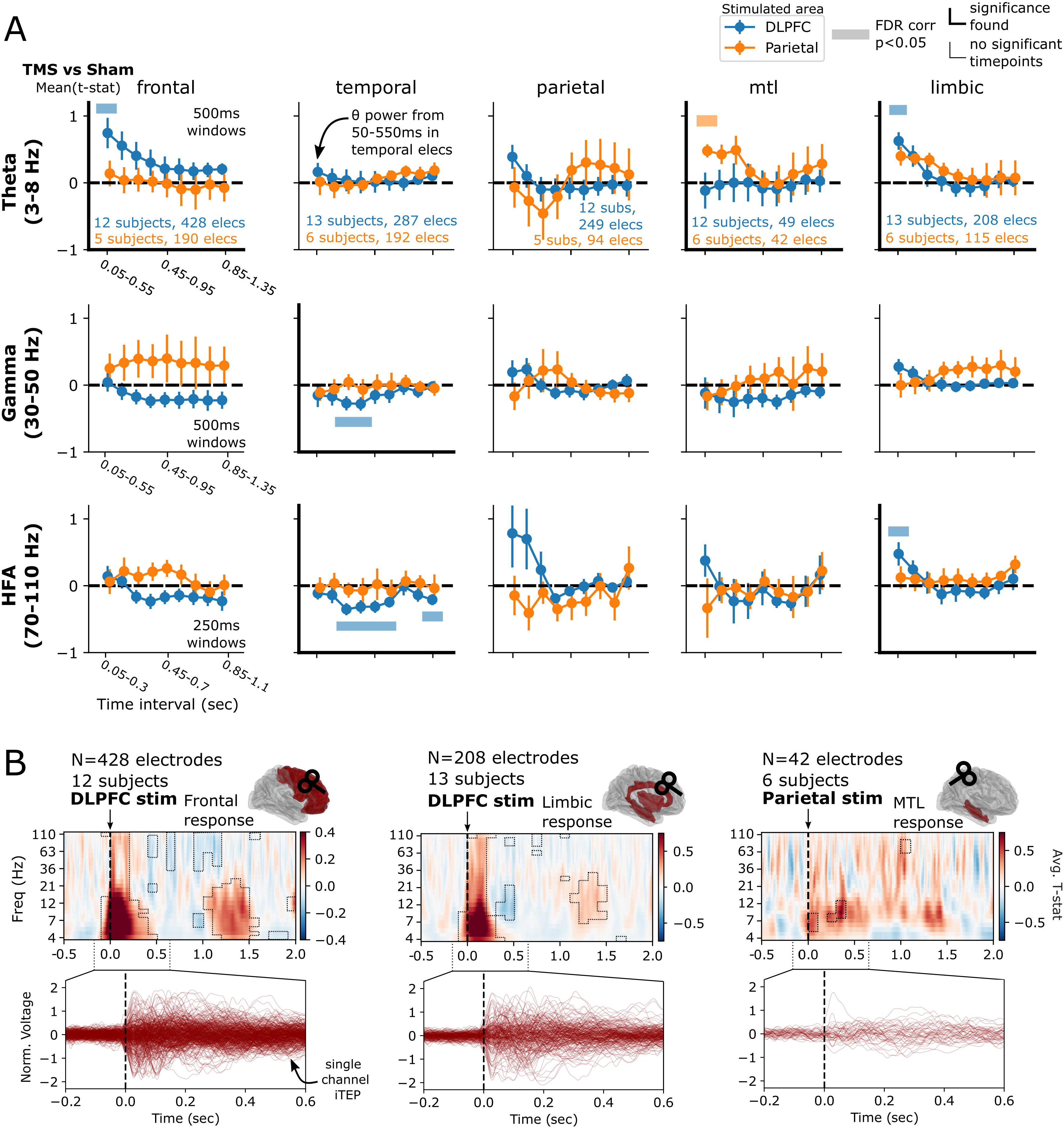
Differential regional responses to DLPFC versus parietal TMS. **(A)** For each frequency band (theta, gamma, and high-frequency activity [HFA]), spectral power is measured in 500ms (theta and gamma) or 250ms (HFA) windows in 100ms steps, for every electrode and subject in the dataset. TMS-related power is statistically compared to sham-related power, as outlined in Figure 1 and *Methods*. The resulting *t*-statistics are averaged across all electrodes within each subject, then across subjects, to generate curves reflecting the change in power over time. Significant differences between TMS and sham-related power are denoted with horizontal bars (*P*<0.05, FDR corrected over timepoints but not regions or frequencies) and are highlighted with thickened axes. In the theta (3-8Hz) band, there was a significant early response in frontal cortices and limbic areas to DLPFC stimulation, while the MTL shows a significant early response to parietal stimulation. Temporal electrodes showed a significant gamma decrease in the intervals starting at 250-350ms, as well as a significant and sustained suppression of high-frequency activity (HFA; 70-110Hz). See **Supplemental Table 1** for a list of structures included in each area. Error bars show +/- 1 standard error of the mean over subjects (SEM). **(B)** *Top:* For each of the three regions showing a significant theta response, time-frequency spectrograms were averaged across all electrodes in each region, demonstrating brief (<500ms duration) increases in theta power immediately following stimulation, with lesser increases seen between 1-1.5 seconds post-stimulation. Boxed areas represent significant TMS vs. sham differences (*P*<0.05, FDR corrected). Time-frequency spectrograms were not generated for regions without a statistically significant theta response. *Bottom*: Butterfly plots representing the trial-averaged voltage response for each electrode across all subjects in a given region (i.e. an intracranially-recorded TEP, or iTEP).

As this global analysis averages across frequency bands and large time windows, it may obscure interesting dynamics that do not cleanly align with predefined time ranges or frequency bands. Within each of the theta-responsive regions identified above, we further analyzed each region by averaging the time-frequency responses across electrodes and subjects, testing each time-frequency “pixel” for a significant TMS vs. sham difference, and correcting for multiple comparisons (**Figure 2B**, *P* < 0.05; see *Methods*). This time-resolved analysis necessarily means stimulation artifact may contaminate power measures close to the pulse, though see *Methods* and *Discussion* for further considerations. Consistent with the previous analysis, these results demonstrated initial broadband power increases to DLPFC stimulation as recorded in frontal and limbic cortices, with the strongest effect in theta and alpha bands (approximately 4-13Hz). DLPFC stimulation suppressed alpha/beta (approximately 9-21Hz) power between 300-500ms post-stimulation in limbic regions. DLPFC stimulation also induced a prolonged (up to 1200ms) gamma and HFA suppression in frontal cortices. Furthermore, after DLPFC stimulation both limbic and frontal regions demonstrated a weaker, later (1-1.5s) increase in theta, alpha, and beta power. Finally, parietal stimulation provoked a significant early (0-400ms post-stimulation) theta and alpha-range increase in the MTL.

Taken together, our results align with prior noninvasive literature which suggests TMS provokes generally widespread and brief responses in lower frequencies (*22*, *24*, *44*). This low-frequency response overlaps considerably with the general spectral profile of non-invasive TMS-evoked potentials (TEPs; see *Separating evoked from induced rhythmic activity* for further analysis, as well as *Discussion*). We extend these findings by (1) demonstrating that such responses are also observed in subcortical areas including MTL and (2) observing a prolonged frontotemporal gamma and HFA suppression, frequencies which are not easily interpreted using scalp EEG. Though only qualitatively compared, we note that neural responses appear to segregate according to stimulation site: DLPFC stimulation provoked low-frequency power in frontolimbic areas, whereas parietal stimulation drove activity within the MTL.

While lobe-level regions-of-interest increase power to detect large-scale (but potentially weak) effects, they obscure the substantial heterogeneity in sub-regional responses to stimulation. Narrowing our view to the lobes and timepoints which demonstrated a significant effect in **Figure 2**, we asked which particular subregions contributed to these effects (using the DKT parcellation, any region with 5 or more subjects sampled). With DLPFC stimulation, early theta power increases (50-550ms) were significant in the precentral, cingulate, and anterolateral frontal regions, particularly orbitofrontal cortex (*P*<0.05 FDR-corrected, **Figure 3A**). Suppression of gamma and HFA power (beginning 250ms following stimulation) in the temporal lobe was significant in the superior and middle temporal gyri (**Figure 3B-C**). Finally, increased limbic HFA power in the 50-300ms interval was localized to the insula and isthmus of the cingulate gyrus (**Figure 3C**). No significant subregions emerged when analyzing the effect of parietal stimulation on MTL theta activity.

**Figure 3.**
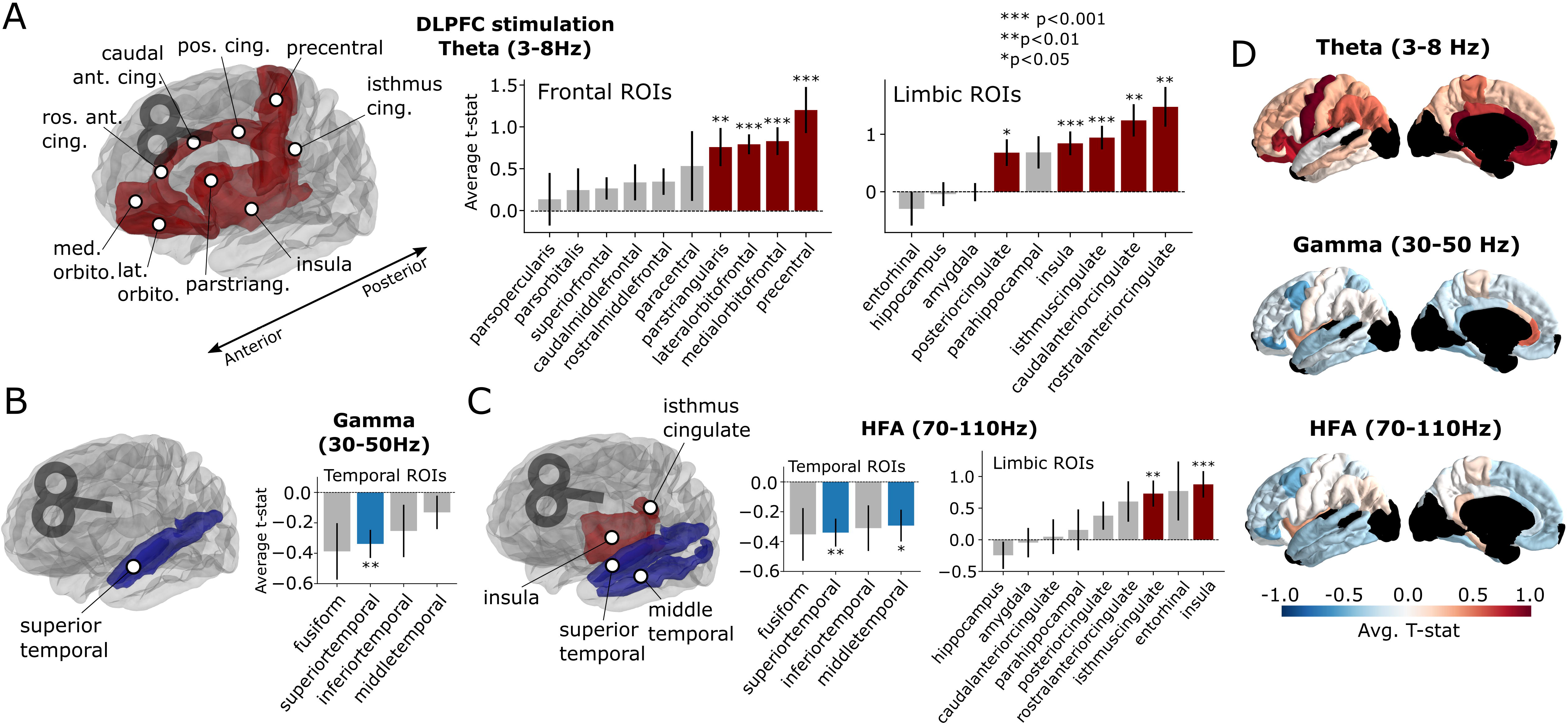
Sub-regional responses to DLPFC stimulation. **(A)** Frontolimbic theta-band responses to DLPFC stimulation. *T-*statistics were computed by comparing responses to DLPFC stimulation between TMS and sham trials in the 50-550ms interval. Time windows were chosen based upon the significance analysis in Figure 2 (see *Methods* for details). DKT regions with significant (*P*<0.05, FDR corrected) TMS-related activity are demarcated in red. **(B)** Temporal lobe gamma-band responses to DLPFC stimulation in the 250-750ms interval, with significant decreases demarcated in blue. **(C)** Temporal lobe HFA decreases following DLPFC stimulation in the 250-500ms interval (left) and limbic increases in the 50-300ms interval (right). **(D)** Representation of average *t*-statistic for each ROI across all subjects and electrodes, in the 50-500ms (theta), 250-750ms (gamma), and 250-500ms (HFA) intervals. Dark regions reflect areas with limited available data (fewer than 4 subjects). See **Supplemental Figure 2** for un-thresholded ROIs and parietal stimulation effects. Error bars reflect +/- 1 standard error of the mean across recording contacts. \**P*<0.05, \*\**P*<0.01, \*\*\**P*<0.001, FDR-corrected across ROIs.

### Separating evoked from induced rhythmic activity

Having established that TMS can alter spectral power in disparate brain regions, we next asked if stimulation also exerts an effect on the *phase* of rhythmic activity. The phase of brain rhythms carries useful information about the nature of a rhythmic signal and may also reveal changes in brain dynamics that were not detectable by measuring changes in spectral power alone. Specifically, when interpreted alongside amplitude increases, the phase consistency of rhythms evoked by TMS helps characterize them as either “induced” or “evoked,” i.e. whether a stimulation pulse provokes rhythms with low or high phase consistency over trials, respectively (*33*). Evoked rhythms, by definition, feature a consistent delay from the stimulation pulse (high phase consistency), while induced rhythms may occur at varying points in time (low phase consistency). In this study, we used the inter-trial phase locking metric (IPL), a commonly-used quantification of phase consistency across trials, computed for each individual electrode (see *Methods* for details). We focused on the theta (3-8Hz) band, which is well-established as containing cognitively and physiologically-relevant phase locking properties, especially relative to higher frequencies (e.g. gamma and higher) (*45*–*47*).

Briefly, IPL was measured by extracting continuous theta-band phase in the post-stimulation time period, and then computing the phase-locking value (i.e. consistency of phase) across all trials for TMS and sham events, separately (**Figure 4A**). In doing so, a continuous measure of the difference between TMS-IPL and Sham-IPL can be computed for each electrode (**Figure 4B**). By measuring the difference between TMS-IPL and Sham-IPL, and then averaging across electrodes into regions-of-interest, we derive a statistical measure of the TMS-related IPL, analogous to the power *t*-statistics shown earlier (**Figure 2**). Following DLPFC stimulation, we found that, across lobe-level ROIs, there was significant TMS-related IPL provoked across several areas, though effects only survived multiple comparisons correction in the frontal and limbic ROIs (**Figure 4C top row**; frontal: Wald *z* = 5.75, *P* < 0.001; limbic: *z* = 5.9, *P* < 0.001). The effect was strongest and significant in the early post-stimulation period (frontal: through the 0.6-1.1 sec interval; limbic: through the 0.5-1 sec interval). In all regions, IPL decayed monotonically over the inter-stimulation period.

**Figure 4.**
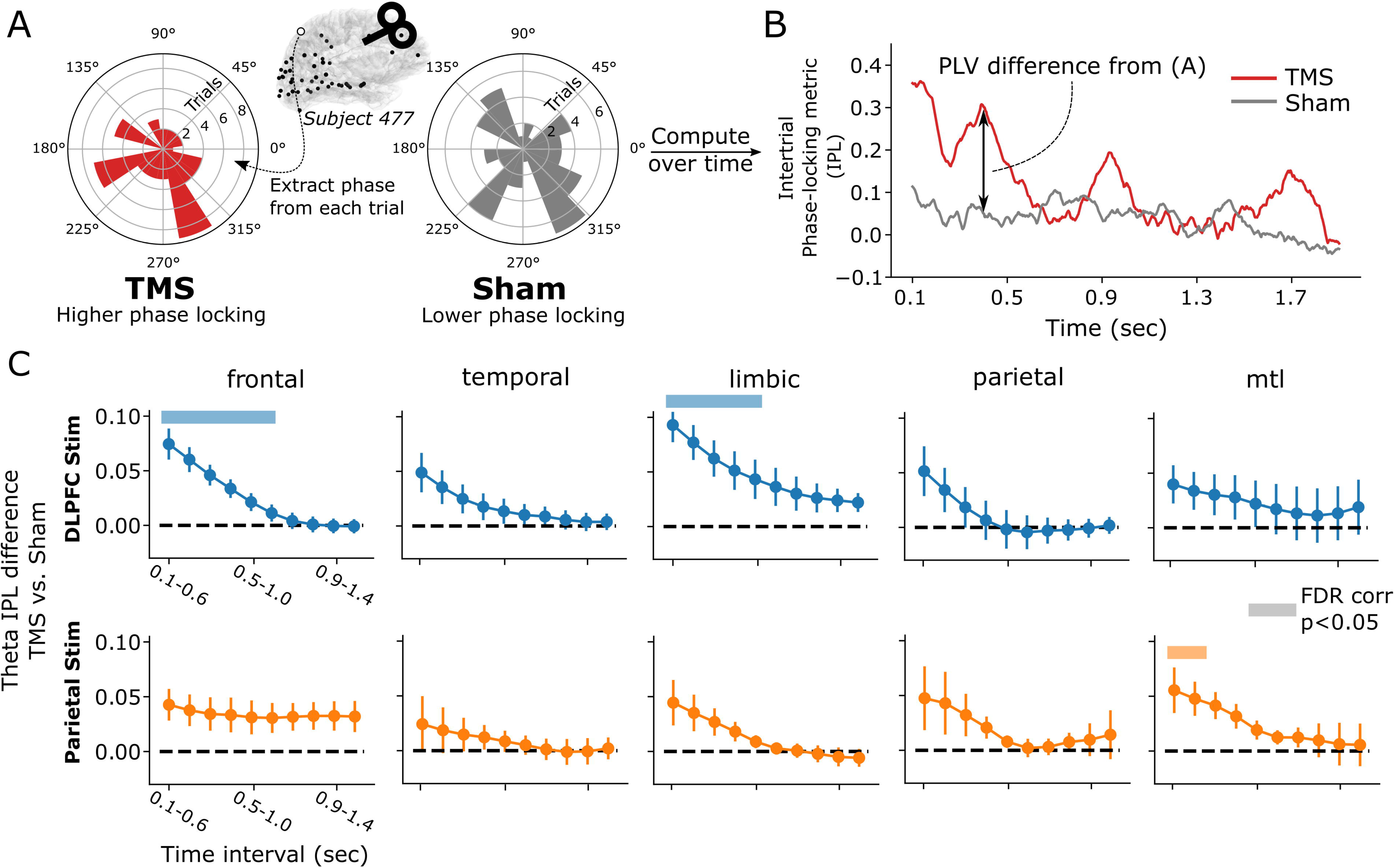
TMS-related theta (3-8 Hz) band inter-trial phase locking. **(A)** Example of inter-trial phase locking value (IPL) computed at one superior parietal electrode from a representative subject, assessed at approximately 400ms after DLPFC stimulation. In this example, there was an elevated inter-trial phase locking in TMS trials – indicating a consistent theta phase across trials – which manifests as a circular phase distribution with clustering in a particular direction (i.e. to one side of the 0 degree axis). Sham trials exhibit a phase distribution with higher variance, corresponding to a lower IPL. **(B)** Timecourse of the IPL for TMS and sham trials from the electrode highlighted in (A). **(C)** Average TMS-minus-sham IPL difference across all subjects and electrodes for the same broad regions used in Figure 2, in 500ms windows spanning the post-stimulation period. Colored bars indicate timepoints when the PLV difference significantly differs from zero (linear mixed effects modeling, FDR corrected *P* < 0.05; see *Methods* for details). Frontal and limbic cortices showed early increases in phase locking in TMS relative to sham trials, following DLFPC stimulation. MTL regions showed a significant early increase driven by parietal stimulation. In all regions, early increases in phase-locking decays towards zero by approximately 1-second following stimulation. Error bars show +/- 1 SEM across subjects. See Figure 2A for the count of subjects and electrodes for each stimulation-region combination.

Following parietal stimulation, effects are similarly early and rapidly decay (**Figure 4C, bottom row**). Significant effects (FDR-corrected *P* < 0.05) were only found in the MTL (through the 0.2-0.7 sec interval; *z* = 2.66, *P* = 0.007), mirroring our results when examining spectral power alone. Similar to DLPFC, these regions showed a steady decay in TMS-minus-sham IPL that tended to approach zero around 500ms post-stimulation.

While elevated theta power and phase-locking in the immediate post-stimulation period would be consistent with the spectral profile of an evoked rhythm, it does not rule-out the possibility that induced oscillations also occur following stimulation. To assess this, we adopted the “fitting oscillations and one-over-f” method (*48*) (FOOOF) to detect theta oscillations. Briefly, FOOOF separates neural power spectra into periodic and aperiodic components, the former of which would be reflective of neural oscillations as opposed to non-oscillatory changes in a signal (*49*). As FOOOF operates on the level of individual trials – and not averages – FOOOF will be sensitive to induced oscillations that occur with variable delay from the stimulation pulse. Conversely, as FOOOF detects circumscribed peaks in the power spectrum, it will be less sensitive to the broadband effects of an evoked potential.

Using FOOOF, we reanalyzed the subset of frontolimbic regions which, following DLPFC stimulation, demonstrated increases in theta-band power using standard spectral methods (these differ slightly from those in **Figure 2A** and **Figure 3A**, as spectral windows were adjusted to better equate to requirements of FOOOF; see *Methods* for details). FOOOF was applied to every stimulation and sham trial, identifying trials in which theta oscillations occurred (**Figure 5A**). The degree to which theta oscillations were observed in TMS was compared to sham trials to generate a new *t*-statistic, based purely on a measure of oscillatory signal. We found that, among the five regions with elevated theta-band power, three (lateral orbitofrontal, medial orbitofrontal, rostral anterior cingulate) demonstrated significant TMS-related increases in oscillatory activity while two became non-significant (precentral gyrus, isthmus of the cingulate; **Figure 5B-C**). Across all DKT regions, regardless of change in power, there was a significant linear correlation between standard spectral power and FOOOF-detected oscillations (*r*(29) = 0.519, *P*=0.004; **Figure 5D**). This suggests that TMS may induce theta oscillations in anterior frontal areas and evoked responses elsewhere.

**Figure 5.**
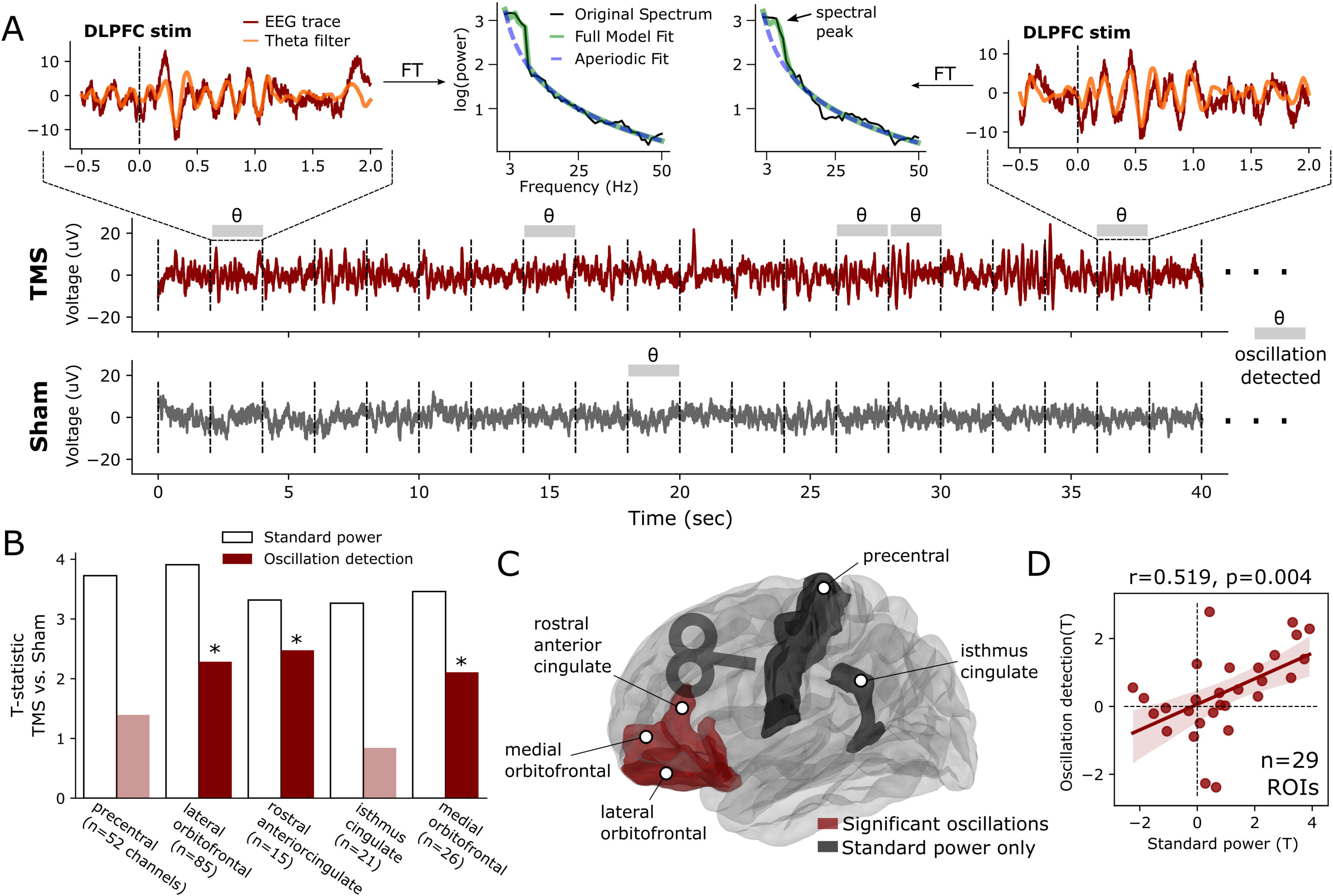
Analysis of theta oscillations provoked by DLPFC TMS. **(A)** Demonstration (n = 1 contact) of FOOOF method for detecting oscillations, applied to a sample trace from an electrode in the medial orbitofrontal cortex. Trials in which a FOOOF-detected theta oscillation occurred are indicated with grey bars. Two example trials are highlighted, demonstrating emergence of a theta oscillation which is reflected by a peak in the spectral power distribution (top center). The theta-filtered signal is overlaid in orange on the original trace. In this example contact, theta oscillations are detected less often in the sham condition. **(B)** TMS-versus-sham *t*-statistics assessed using standard power (white bars) or FOOOF oscillation detection (red bars). Regions with significant (*P*<0.05) oscillatory modulation as detected by FOOOF are indicated with asterisks and darker color. **(C)** ROIs which demonstrated significant TMS-related modulation of theta oscillations using FOOOF (red) and those in which raw spectral power increased but theta oscillations were not found (black). **(D)** Overall correlation between TMS-related oscillations and raw spectral power, across all available ROIs regardless of significance. There is a significant (*r*(29) = 0.519, *p*=0.004) positive linear correlation.

### Subcortical responses to cortical TMS

Subcortical structures play a key role in neuropsychiatric illness, but non-invasive electrophysiological recordings such as scalp EEG are unable to accurately localize spectral activity from subcortical areas. Combined TMS-iEEG offers a unique opportunity to directly record from subcortical areas during concurrent TMS. In the previous analysis, we broadly demonstrated a low-frequency response to single pulse TMS in the MTL and limbic areas, across large time windows and frequency bands (**Figure 2**). We now examine the specific spectral dynamics which emerged in the hippocampus and amygdala – two areas which are (1) sufficiently sampled in our cohort (i.e. at least 5 subjects) and (2) strongly implicated in neuropsychiatric illnesses(*50*–*52*). Our purpose is to characterize the full spectro-temporal responses to cortically-targeted TMS in these functionally and anatomically distinct structures. In doing so, we hope to shed light on how propagated neural activity from the cortex manifests as subcortical rhythmic activity.

Averaged across all hippocampal recording contacts and subjects, DLPFC stimulation suppressed activity in the gamma band between 400 and 500ms post-stimulation (**Figure 6A**; Wald *z* = -3.4, *P* < 0.001) and HFA band between 400 and 600ms post-stimulation (*z* = -3.3, *P* < 0.001). (Nonsignificant increases in low-frequency power were appreciable principally in alpha and theta bands.) No significant high-frequency suppression was observed after hippocampally-targeted parietal stimulation. However, a subthreshold increase in theta power (*P* < 0.05 uncorrected; maximum of Wald *z* = 2.1 between 300-400ms) is seen in the first 500ms following parietal stimulation, which likely contributed to our earlier finding of theta power increases in the MTL more broadly. In the amygdala, a significant decrease in theta power was seen 400-500ms after DLPFC stimulation offset (*z* = -3.5, *P* < 0.001), alongside subthreshold early increases and later decreases in broadband spectral power. Parietal stimulation results in no significant spectral modulation in any band after correction for multiple comparisons (**Figure 6B**). Broadband amygdala response after DLPFC stimulation raised the question of possible stimulation artifact contamination. By examining the spectral responses of individual electrodes, the broadband-appearing response was rather driven by substantial inter-subject and inter-electrode variability in peak responses, which manifests as a broadband response in the statistical average – without evidence for contamination by stimulation artifact (**Supplemental Figure 3**).

**Figure 6.**
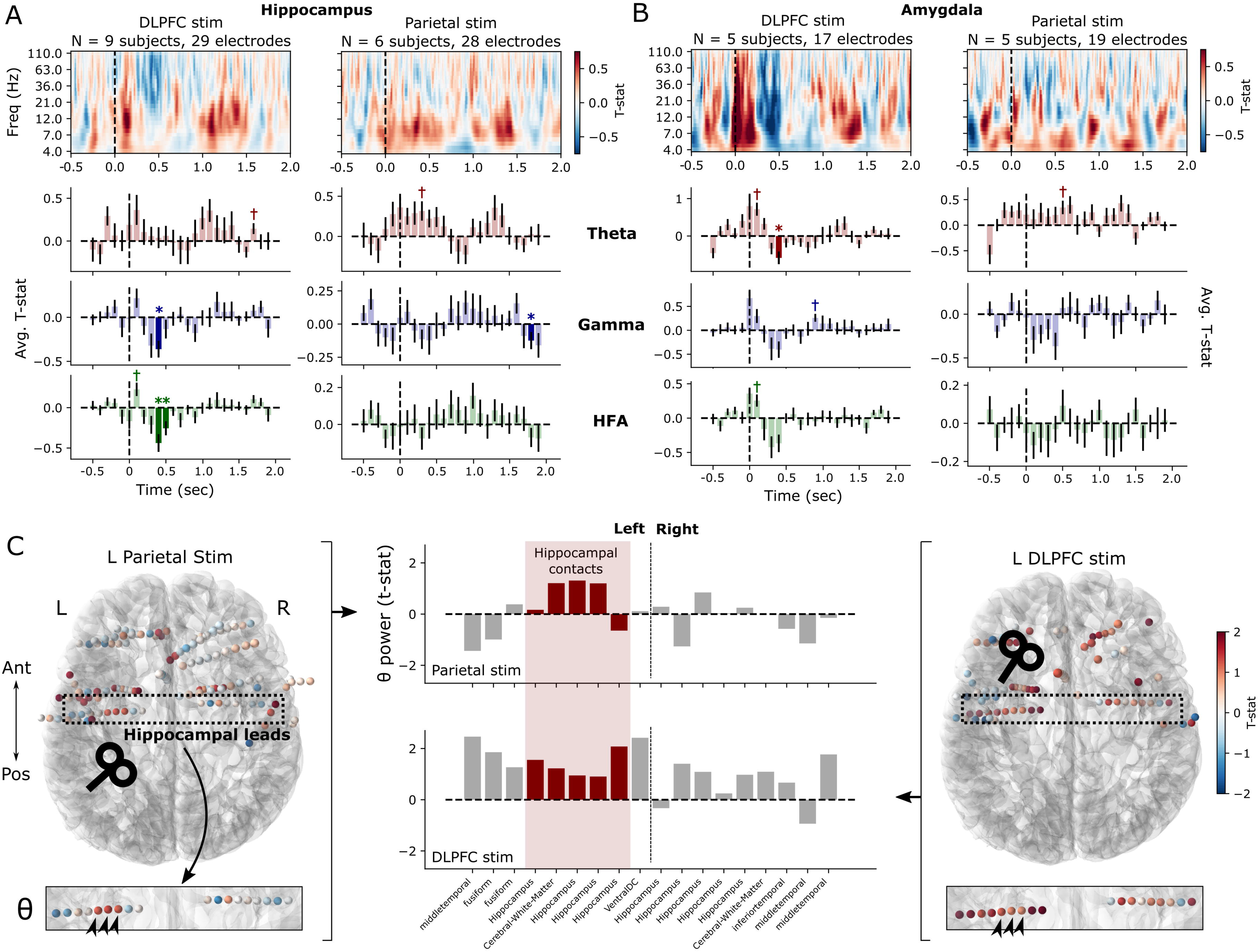
Subcortical responses to TMS in hippocampus and amygdala. **(A)** *Top:* Time-frequency spectrograms of the hippocampal response to DLPFC and parietal stimulation targeted towards the cortical area with maximal functional connectivity to the hippocampus. These reflect the average TMS-minus-sham difference across all electrodes and subjects; no statistical corrections are applied. Dotted line indicates time of stimulation pulse. *Bottom:* Timecourses of band-averaged power in the theta, gamma, and HFA bands, with indicators for significant TMS-minus-sham differences in 100ms windows (\**P*<0.05, FDR corrected across timepoints, ✝*P*<0.05 uncorrected; see *Methods* for details). Error bars show +/- 1 SEM across contacts. **(B)** As in (A), but for amygdala spectral responses to DLPFC and parietal stimulation. **(C)** In a single subject who underwent both DLPFC and parietal stimulation, each recording contact is colored by the average theta power following TMS vs. sham stimulation (50-550ms post-stimulation). The hippocampal electrode leads were specifically examined (dotted box) to understand the differential responses in this structure to cortical stimulation. Theta power is represented for each recording contact along these leads (center bar plots) with hippocampal contacts highlighted in red. Three recording contacts (arrowheads) show a qualitatively increased response from parietal stimulation (left) which is less anatomically specific following DLPFC stimulation (right).

Our use of hippocampally-targeted parietal stimulation – via resting-state fMRI functional connectivity – raises the question as to whether such targeted stimulation yields specific hippocampal responses. In general, as shown earlier, the MTL exhibited a significant low-frequency response to parietal stimulation which was not evident in DLPFC stimulation (**Figure 2A**). And on average, the hippocampus itself demonstrated a subthreshold increase in theta power following parietal, but not DLPFC stimulation (**Figure 6A**). Given the underlying heterogeneity of responses to TMS, we asked whether specific hippocampal responses were evident at the single-subject level (**Figure 6C**). Only two subjects (with hippocampal contacts) received both DLPFC and parietal stimulation; one of those two demonstrated a qualitatively specific increase in hippocampal theta power in response to parietal stimulation, while exhibiting broad, nonspecific theta increases following DLPFC stimulation (which were lesser in magnitude in the hippocampus relative to ipsilateral cortical areas). The other subject did not demonstrate a qualitatively or quantitatively specific response in the hippocampus following parietal stimulation.

The interplay between DLPFC and anterior cingulate cortex (ACC) is an area of significant interest. Prior evidence indicates that TMS can be used to alter ACC activity and connectivity (*37*, *53*). Given this background, we further analyzed the spectral response to DLPFC-targeted TMS in cingulate subregions (**Figure 7**). The ACC as a whole showed a brief significant response in theta, gamma, and HFA bands from 100-200ms after stimulation (*P* < 0.05, FDR corrected). Upon inspecting ACC subregions, we found that the rostral ACC demonstrated a robust and statistically significant increase in theta power following TMS, persisting until 600ms post-stimulation. The caudal ACC showed no significant effect during that interval in theta, though continued to demonstrate significant early (100-200ms) increases in gamma and HFA. Posterior cingulate cortex (PCC) demonstrated no significant spectral modulation in any prespecified frequency band. (We qualitatively noted a suppression of alpha and low-beta power, approximately 9-15Hz, in the 250-500ms interval.) Of note, fewer than 5 subjects had parietally-targeted TMS and cingulate recording contacts, precluding analysis of cingulate response to parietal TMS.

**Figure 7.**
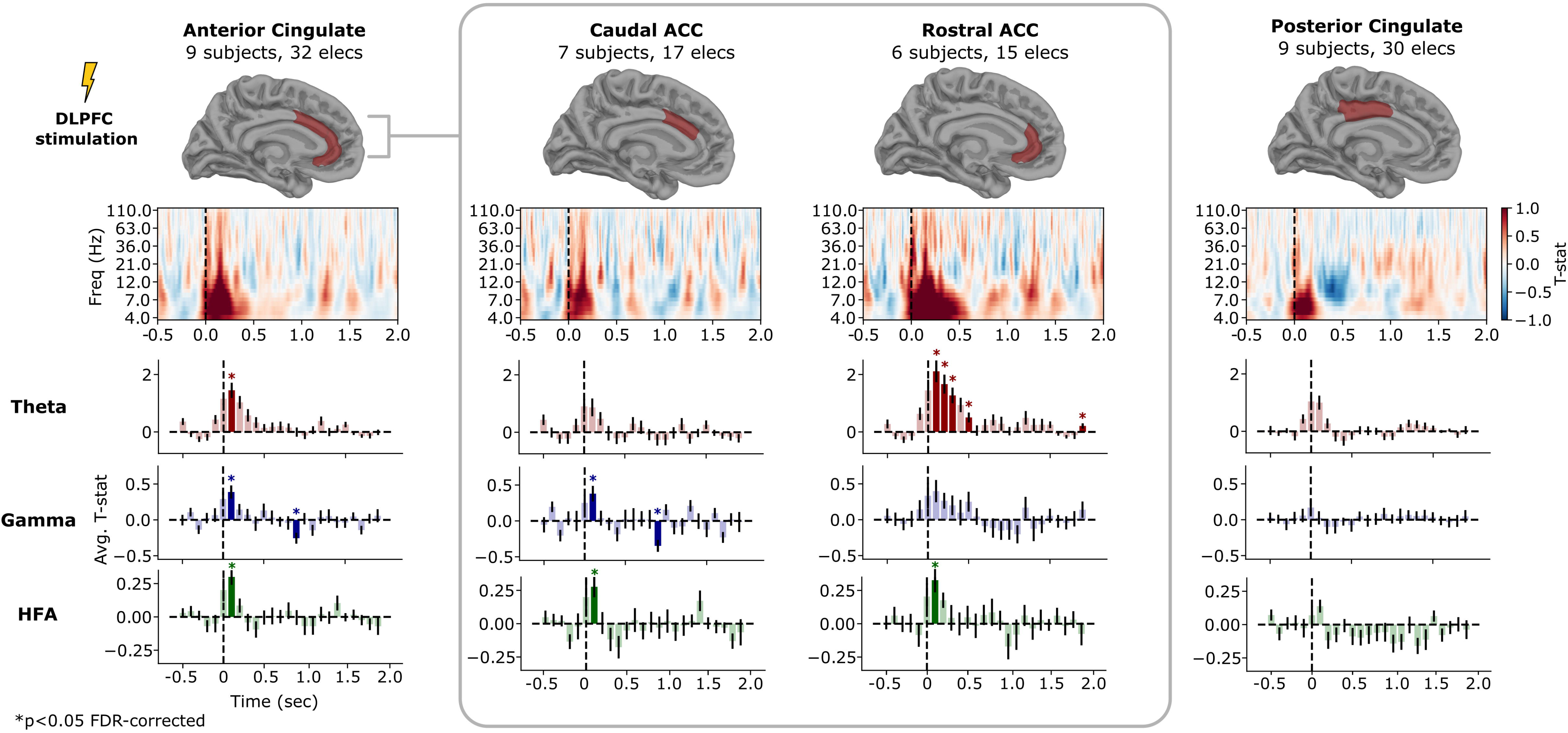
TMS-related spectral power within cingulate cortex following DLPFC stimulation. Analysis conducted as described in Figure 6. Anterior cingulate (ACC) demonstrates statistically significant TMS-related power in the 100-200ms interval following DLPFC stimulation in all three frequency bands assessed; there was a more prolonged theta-band effect in underlying rostral ACC with a significant TMS-minus-sham difference extending to the 500-600ms post-stimulation interval. Caudal ACC demonstrated no theta effect but did show a 100-200ms increase in gamma and HFA power. Posterior cingulate demonstrated no significant spectral modulation in any prespecified frequency band, but did exhibit a qualitative increase early increase in theta power and approx. 200-500ms decrease in alpha power, on inspection of the time-frequency response. Significance tests conducted using a linear mixed-effects model as described in *Methods*; error bars show +/- 1 SEM across recording contacts. \**p* < 0.05, FDR-corrected over timepoints.

Taken together, these analyses of subcortical responses in small samples or individual subjects should be interpreted in their statistical context: early hints of the possibility that TMS can be used to modulate the spectral activity within subcortical structures. Specifically, there is a statistical finding that DLPFC stimulation can suppress high-frequency activity in the hippocampus and low-frequency activity in the amygdala. No statistically significant effects emerge following parietal stimulation, but we found a notable increase in hippocampal theta power within 500ms of stimulation offset that does not reach corrected significance. As these sample sizes are small and may exhibit substantial inter-individual variability, further work is needed to validate whether this evidence is idiosyncratic, or reflects a repeatable means to non-invasively and predictably modulate subcortical rhythmic activity.

## Discussion

TMS is used to modulate neural circuits in neuropsychiatric illness, but until recently, human brain responses to stimulation could only be understood by non-invasive, spatiotemporally imprecise methods. In this study, we used indwelling electrodes to measure intracranial responses to TMS, allowing for signals that are more precise and have higher spatiotemporal resolution that can be decomposed in the full spectral domain. In doing so, we aimed to (1) characterize the spectral responses of key brain regions to TMS, further asking whether induced oscillations can be differentiated from evoked rhythms, (2) assess responses in higher frequency bands that have been difficult to interpret using scalp EEG, and (3) examine the responses of deep brain structures to TMS, shedding light on how downstream regions respond to propagated activity from stimulated cortex.

We found that DLPFC stimulation tended to cause brief, early increases in theta power in frontal and limbic cortices, particularly the orbitofrontal and anterior cingulate cortex (ACC; **Figure 3A** and **Figure 7**). Higher frequencies, including gamma and HFA bands, were suppressed predominantly in the temporal lobe, though smaller effects were also noted frontally. Parietal stimulation – which was targeted based on functional connectivity to the hippocampus –caused significant early theta increases in the MTL, but in no other regions or frequency bands. In the hippocampus, DLPFC stimulation attenuated high-frequency activity, with indications that parietal stimulation may increase low-frequency activity. Phase analysis suggested that the frontolimbic low-frequency increases were driven by an evoked and phase-locked response to stimulation, but a trial-level analysis confirmed the co-occuring presence of induced theta oscillations within anterior frontal regions.

Taken together, these results demonstrate that TMS provokes brain-wide changes in spectral power across frequency bands, consistent with prior evidence from combined TMS and scalp-EEG (*7*, *26*, *27*). We extend this work in several important ways. First, we established that these power increases reflect a mixture of evoked rhythms and induced theta oscillations that may be region-specific. The relative contributions of these two neural processes are likely not equal; the strong phase-locking we observed alongside early and typically brief increases in spectral power (i.e. less than 500ms) is a good indicator that an evoked response is the predominant driver of increases in low-frequency power. Traditional spectral methods, such as measurement of trial-averaged spectral power, would be unable to disentangle the two phenomena as both feature low-frequency sinusoidal fluctuations in the local field potential. As such, it was critical to implement analytic tools that specifically look for circumscribed peaks in the power spectrum (*48*, *49*, *54*).

Second, our study provides a view into high-frequency activity that is beyond what can be offered by scalp EEG (*20*, *55*, *56*). Moreso than the low-frequency increases, early (approximately 200-500ms) decreases in gamma and HFA were consistent and statistically significant in widespread regions following DLPFC stimulation, especially in temporal cortices (including medial temporal regions). These frequency bands have been hypothesized as signatures of population neural spiking, particularly in high gamma and HFA (*42*). Accordingly – because HFA likely relates to neural firing – our findings suggest TMS to the DLPFC may suppress neural firing. This effect may be either directly related to the immediate neural effect of the TMS pulse or indirectly as an aftereffect of the cortical silent period that often occurs after the early response in an evoked potential. The effects of TMS on hippocampal HFA and gamma signatures are particularly notable, as these deep neural responses could not be reliably elucidated with non-invasive measures or without the high temporal resolution of iEEG.

Future work should ask whether alternative stimulation paradigms – such as repetitive or rhythmic (e.g. theta-burst) stimulation – would be more likely to provoke induced as opposed to evoked rhythmic activity. Prior work examining the cortical responses to patterned direct intracranial stimulation suggest this would be the case by demonstrating prolonged power increases or non-timelocked events (*17*, *20*, *56*–*58*), though this remains to be established with TMS. Our finding of high-frequency suppression could be explored further with measures of single-unit activity, explicitly linking the effect of TMS to neural firing. Additionally, the effect of TMS on high-frequency activity may reflect the relationship between the onset of a pulse and the phase of ongoing background oscillations, which can either be explored analytically or in closed-loop designs.

Functionally-targeted parietal stimulation provoked qualitatively greater responses in the MTL as compared to frontal stimulation (**Figure 2A, 6C**), though we had insufficient data to statistically measure this effect within-subjects. This is suggestive that functional connections dictate the way in which stimulation propagates through the brain, extending a growing body of literature (*16*, *18*, *59*, *60*). However, further work is needed to determine if this principle generalizes to other stimulation sites and recording areas, and more data is needed to extend this finding beyond qualitative observation. Moreover, as our focus was to characterize modulations of region-specific power and phase locking, we did not ascertain whether individual variability in functional connectivity correlates with the effect of stimulation in downstream regions. To answer this question, we plan to analyze the relationship between subject-specific measures of intracranial functional connectivity (e.g. resting-state fMRI or electrophysiological coherence) and the TMS-provoked activity recorded at indwelling electrodes.

From these data, it is also clear that non-invasive, cortically-targeted stimulation can modulate electrical activity in deep brain structures that are not directly accessed by stimulation itself, extending previous work in direct cortical stimulation (*61*, *62*) and fMRI (*11*, *63*). Within the limits of the moderately-sized samples in this study (up to 17 subjects depending on the stimulation site and recording location), our data suggest that TMS directed at the DLPFC – at least following single pulses – suppresses high-frequency activity in the hippocampus for several hundred milliseconds. The ability to predictably suppress neural firing in the hippocampus with cortically-targeted TMS could point towards its therapeutic potential, especially in psychiatric illness that features pathological hippocampal activity, such as depression (*64*) or psychosis (*65*). There is also weak but intriguing evidence that parietal stimulation can instigate theta rhythms in the hippocampus – a finding that has profound implications for how we might use stimulation in modulating core cognitive functions of the hippocampus itself (*43*, *66*, *67*).

Although the mapping between stimulation and amygdala responses is less clear, these results highlight the potential for using TMS to precisely modulate the function of deep brain structures with profound implication in neuropsychiatric disease. Future work should focus on clarifying how cortically-propagated signals exert their effects on these structures, and whether specific stimulation patterns can be used to provoke specific frequency responses (*68*).

Stimulation-response paradigms often raise the question of spectral contamination by the pulse artifact. While it is difficult to ensure that zero artifactual components enter these kinds of analyses, several features of our analytic methods and results suggest the effect would be small. First, our core analyses used windowed measures of spectral power that contain no data from the stimulation period itself, beginning 50ms after stimulation offset. To subsequently provide a fuller spectral representation of stimulation’s effects in specific areas, time-frequency representations in **Figure 2C**, **Figure 6A-B**, and **Figure 7** necessarily do overlap with the stimulation interval. However, care was taken to remove and re-interpolate the stimulation artifact period, as described in *Methods* – our demonstration of peak effects more than 100ms after stimulation suggests that this data cleaning was successful. Finally, we demonstrated decreases in high-frequency power starting several hundred milliseconds after stimulation (e.g. **Figure 6A**), which would be very unlikely to occur due to stimulation artifact alone.

These findings represent a key advance in how we understand the mechanisms of TMS-related change in neural function. More broadly, they demonstrate the promise of combining non-invasive stimulation with direct intracranial recordings, providing a window into the detailed electrophysiology of brain stimulation while developing a therapeutic technique that can be easily deployed in outpatient clinical settings.

## Materials and Methods

### Human subjects

Seventeen neurosurgical patients with medically intractable epilepsy underwent a surgical procedure to implant intracranial recording contacts on the cortical surface (electrocorticography) and within brain parenchyma (stereo-EEG). Contacts were placed in accordance with clinical need to localize epileptic regions. Each patient was admitted to the University of Iowa Hospitals and Clinics for 14 days of clinical and electrophysiological monitoring to identify their seizure focus. TMS experiments were conducted after the final surgical treatment plan was agreed upon between the clinical team and the patient, typically 1-2 days before the planned electrode explantation operation and 24 hours after the patient had restarted anti-epileptic medications. All experimental procedures were approved by the University of Iowa Institutional Review Board, who reviewed safety data from a separate experiment prior to approval for human subjects (*37*). Written informed consent was obtained from all participants.

### Imaging protocol and intracranial electrode localization

Intracranial electrodes were localized in a manner identical to that described in Wang, et al. (2022)(*37*). Briefly, patients underwent anatomical and functional MRI scans within two weeks of electrode implantation, including resting-state functional MRI (rsfMRI). The day following implantation, subjects underwent a second MRI and volumetric computerized tomography (CT) scans. The location of each contact was identified on the post-implantation T1-weighted MRI and CT, and subsequently post-implantation scans were transformed to pre-implantation T1 anatomical space in a manner that accounts for the possibility of post-surgical brain shift (*69*). Freesurfer (*70*) was used to map electrode locations onto a standardized set of coordinates across subjects, which were then labeled according to their location within the Desikan-Killiany-Tourville (DKT) anatomical atlas.

### Transcranial magnetic stimulation

For stimulation, we used a MagVenture MagVita X100 230V system with a figure-of-eight liquid-cooled Cool-B65 A/P coil (Magventure; Alpharetta, GA, USA). Stimulation pulses were biphasic sinusoidals with a pulse width of 290 microseconds, with stimulator output set at a percentage of each subject’s motor threshold. Pulses were delivered at 0.5Hz, allowing for 2-second inter-stimulation intervals to examine spectral responses. TMS experiments were conducted 12-13 days after implantation and after starting antiepileptic medications. Neuronavigation using frameless stereotaxy was guided with Brainsight software supplied with the pre-implantation T1/MPRAGE anatomical scan. Stimulation parameters were recorded in Brainsight during all experimental trials. Motor thresholds were determined starting with the hand knob of the motor cortex as a target, beginning at 30% machine output and adjusted in 5-10% increments until movements were observed in 50% of trials.

In the main experiment, single pulses were directed at DLPFC or parietal targets at or above motor threshold (100% was used if 120% was not tolerated due to pain). DLPFC targets were defined by the Beam F3 region (*71*), identified by transforming published coordinates (MNI 1mm: -41.5, 41.1, 33.4) (*72*) into each subject’s native T1 and displaying it in Brainsight. The stimulation site was adjusted slightly if access was impeded by head wrap or anchor bolts for securing electrodes. Parietal targets were identified by localizing the site within the inferior parietal lobe with maximal resting-state fMRI-based connectivity to the hippocampus. A 4mm spherical ROI was placed at the contact location in the hippocampus to serve as the seed. Mean timecourse in the ROI was calculated, and then a Pearson’s correlation against this timecourse was calculated for every voxel to generate a simple network map. This correlation map was then loaded into Brainsight and thresholded to identify the highest correlation in the lateral parietal cortex posterior to the post-central gyrus, visually confirmed to be the peak correlated voxel nearest to the lateral parietal ROI published in Nilakantan et al. (2019)(*73*).

Sham pulses were delivered in an identical manner to active, with the TMS coil flipped 180 degrees such that the magnetic field was directed away from the head. Participants underwent at least 50 stimulation pulses (“trials”) and 50 sham pulses, though we included subjects with as many as 150 stimulation and 300 sham events, if time and clinical constraints allowed. In one subject, only 33 single pulse TMS trials were included due to tolerability.

### iEEG recording

Electrode recordings were conducted in a manner identical to Wang, et al. (2022)(*37*). Briefly, depth and grid electrodes (Ad-Tech Medical; Racine, WI, USA) were either stereotactically implanted or placed on the cortical surface, respectively. A platinum-iridium strip electrode placed in the midline subgaleal space was used as a reference. Data were amplified, filtered (ATLAS, Neuralynx, Bozeman MT; 0.7-800 Hz bandpass), and digitized at 8000Hz. In all subjects, contacts were excluded from analysis if they were determined to be involved in the generation or early propagation of seizures (412/3894 contacts; 10.6%), if stimulation artifact saturated the amplifier (835/3894; 21.4%), or if electrodes were contaminated by nonneural noise indicative of poor connection or placement outside the brain (67/3894; 1.7%).

### iEEG preprocessing and analysis

The FieldTrip MATLAB toolbox (*74*) was used to load iEEG data into our analysis pipeline. Data preprocessing analysis was principally done with the MNE toolbox (*75*) in Python. First, raw signals were re-referenced to account for large-scale noise or contamination of the reference electrodes; stereo-EEG (depth) electrodes were re-referenced using a bipolar montage, while grids and strips on the cortical surface were collectively re-referenced to their common average.

Although we generally avoided performing spectral analysis on the period of time containing the ∼15ms stimulation artifact itself, some analyses (including generation of time-frequency representations in **Figure 2** and **Figure 6-7**) necessitate analysis of the full interval during each trial. For this reason, we scrubbed the stimulation artifact from all signals and replaced it with synthesized stationary iEEG that reflects a similar spectral profile as the background(*76*). Specifically, the iEEG signal was clipped from 25 ms prior to 25 following stimulation and replaced with a weighted average of the 50ms immediately following and prior to stimulation. Specifically, the pre- and post-stimulation clips were first reversed, then tapered linearly to zero along the length of the signal, and then finally summed together to replace the artifact period. Finally, signals were notch filtered at 60 Hz and harmonics to remove line noise, using an F-test to find and remove sinusoidal components (*77*). Lastly, signals were downsampled to 500 Hz for further analysis.

Our general analytic strategy was to statistically compare spectral activity in TMS trials against sham trials, in order to control for auditory and expectancy effects associated with the stimulation click. To do this, iEEG signals for each contact were segmented into 2.5-second intervals, spanning 500ms prior to stimulation until 2 seconds following stimulation (**Figure 1B**). To first examine broad effects in the large ROIs used in **Figure 2**, we used the multitaper method (time-bandwidth product of 4, excluding tapers with <90% spectral concentration) to measure the power spectral density (PSD) from 3Hz to 110Hz in discrete 500ms or 250ms windows, depending on the frequency of interest. 500ms windows were used in the theta (3-8 Hz) and gamma (30-50 Hz) ranges, while 250ms windows were utilized for high-frequency activity (HFA; 70-110Hz). These differential widths account for the fact that high-frequency activity tends to fluctuate at faster timescales than power in lower frequency bands, making it more appropriate to analyze in briefer time windows. Power was estimated starting 50ms after stimulation to avoid residual contamination from stimulation artifact, in successive overlapping windows spaced 100ms apart until 850ms following stimulation.

### Power responses in large ROIs, subregions, and subcortical areas

In each time window and for each frequency band, powers were log-transformed and averaged over constituent frequencies within the band. To account for drifts in baseline power over time, we subtracted “baseline” power as measured in a 450ms window preceding each stimulation event, buffered by a 50ms gap from the stimulation artifact to avoid any chance of contamination (**Figure 1B**). Baseline power was otherwise measured exactly as per spectral methods described above (*iEEG preprocessing and analysis*). Baseline-corrected powers were compared between TMS trials and sham trials using a two sample *t*-test. This process generated a *t*-statistic for each recording contact in the dataset, at each timepoint and frequency band of interest. Finally, *t-*statistics were averaged across all contacts that fell within a given ROI, for every subject. We did not analyze any region with less than 5 subjects’ worth of data for a given stimulation target.

To generate **Figure 2A**, *t-*statistics were averaged across subjects and tested against zero for significance. Due to the hierarchical nature of our data, variable number of electrodes in each subject, and the possibility of correlated responses between electrodes within subject, we adopted a linear mixed modeling approach (LMM) for major statistical analyses in this manuscript. Specifically, we used the LMM implementation in the Python statsmodels package (*78*). Here, we used intercept-only LMMs to model the variability of *t*-statistics across recording contacts and subjects, specifying subjects as random effects. We used the Wald test to assess the significance of the intercept, asking whether power *t*-statistics significantly differed from zero in our population. Resulting *p*-values were FDR corrected for multiple comparisons over timepoints (α = 0.05). No other effects were included in the model. Note that, in **Figure 2A**, error bars reflect +/- 1 standard error the mean (SEM) over subjects, as the hierarchical variability discussed above cannot be easily graphically represented.

To generate the time-frequency spectrograms in **Figure 2B** and **Figures 6-7**, we slightly modified our analytic approach to allow for the continuous measurement of spectral power, as opposed to discrete windows (**Figure 1B, 2A**). For each contact, we used the Morlet wavelet convolution (3 cycles in length) to extract a continuous measure of power at 25 log-spaced frequencies between 3Hz and 110Hz, log-transformed the result, and subtracted baseline power in the manner described above. We used a 2-sample *t-*test to compare powers between TMS and sham trials, at each pixel of the time-frequency representation. To test for statistical significance of regions within the time-frequency representation (**Figure 2B**), *t*-statistics for each contact in a given ROI were first averaged within subjects, and then tested against zero using 1-sample *t*-tests to generate a *p*-value for each pixel; finally, *p*-values were FDR corrected for multiple comparisons (α = 0.05) to identify time-frequency areas where TMS-related neural activity significantly different from sham activity.

We note that, in using a continuous measure of power over the entire trial, these time-frequency representations may reflect contamination from stimulation artifact, despite efforts to reduce this effect (see *iEEG preprocessing and analysis*). For this reason, our primary statistical analyses were performed on windowed intervals that strictly avoid samples which could contain stimulation artifact (**Figure 2A**, **Figure 3**).

To ascertain the subset of DKT regions which contributed to lobe-level effects (**Figure 3**), we used a series of 1-sample *t*-tests across all recording contacts which fell within a specified region, asking whether there was a significant TMS-related effect in our population. Only regions with sampling from 5 or more subjects were included. Due to the smaller number of data points in these subregions, linear-mixed effects models generally failed to converge, necessitating the use of *t*-tests. These tests were only performed for the lobes and time windows in which a significant effect had been detected (**Figure 2**), constraining the total number of tests. *P*-values were corrected for multiple comparisons.

Analytic methods to measure spectral power in the hippocampus and amygdala (**Figure 6A-B**) were generally identical to those described above. However, to quantify the specific temporal dynamics of how spectral power evolved in these regions after the TMS pulse, we avoided the 500ms windows used to assess large-scale power dynamics as in **Figure 2**. Instead, we first performed Morlet wavelet convolution and then averaged resulting powers into successive non-overlapping 100ms windows, for each frequency band. Statistical testing for significant differences between TMS and sham trials was performed using the same LMM approach as outlined above. Given the smaller number of subjects and electrodes which contributed to these regions, we did not perform statistical testing on the time-frequency representations themselves.

### Inter-trial phase locking (IPL)

To assess the effect of TMS on low-frequency phase locking, we adopted the inter-trial phase locking (IPL) metric, otherwise known as the phase-locking value (*79*). This metric reflects the consistency of phase values, at a given frequency and timepoint, across all trials. High IPL would be indicative of rhythms that are significantly phase-locked to the stimulation pulse, whereas low IPL cannot be concretely interpreted (either reflecting low amplitude rhythms, non-phase locked rhythms, or some combination of both). As for our initial power computations, we again used the multitaper method (time-bandwidth product of 4, two cycles in length, spanning 3-8Hz) as implemented in MNE Python (“tfr_array_multitaper”), which computes IPL by first extracting a continuous measure of phase, and then measures the inter-trial consistency by measuring the mean resultant vector length of phase values across trials. Resulting IPLs fall between 0 and 1, with 1 indicating perfectly consistent phases across trials, and 0 indicating phase distributed uniformly from 0 to 360 degrees.

Since phase-locking is biased by the number of trials that contribute to its computation (*80*), we randomly selected *n* trials from the TMS and sham events in each subject, where *n* is the lower number of trials between the two blocks. In this way, trial counts were matched across TMS and sham events, removing the possibility of PLV bias. IPL was measured starting 100ms after stimulation. As IPL is sensitive to edge artifact, we applied a 450ms “mirror” buffer to the edges of the signal before convolution by reversing the leading and trailing edges of the signal. These buffers were then clipped from the resulting IPL trace prior to further analysis. Finally, as in our power analyses, we measured average IPL In the 450ms “baseline” period prior to each stimulation event (see *Power responses in large ROIs and subcortical areas*), and subtracted this value from post-stimulation IPL for each trial. To measure the TMS-related IPL relative to sham-related IPL, we subtracted the (baseline-corrected) sham IPL from TMS IPL to generate a difference measure (ΔIPL), where positive values would reflect TMS-related increases in inter-trial phase locking.

As in our power analyses, we measured the population effect of TMS on IPL (**Figure 4**) by first averaging ΔIPL for each in 500ms windows spanning the post-stimulation period, beginning at 100ms following stimulation and ending at 900ms in 100ms steps. Next, we averaged ΔIPLs across all contacts within a given ROI, and finally averaged across subjects. As described previously, we used an LMM approach to test the significance of ΔIPL in our population, specifying subjects as random effects in an intercept-only model. *P*-values determined via a Wald test for significance were FDR corrected for multiple comparisons (α = 0.05) (**Figure 4**).

### FOOOF analysis of theta oscillations

To ascertain if specific subregions exhibited induced oscillations, as opposed to evoked low-frequency responses, we relied on the FOOOF toolbox (*48*). By parameterizing the trial-level power spectra into periodic and aperiodic components, FOOOF detects the presence of discrete peaks in the power spectrum which are thought to more faithfully and specifically represent neural oscillations (*49*), as opposed to other phenomena which may alter the spectral profile of EEG.

FOOOF was applied on a per-trial basis to EEG traces from all TMS and sham events (**Figure 5A**), for each subject and electrode in the dataset. We narrowed our view to the subset of regions which exhibited TMS-related increases in raw theta power (**Figure 3A**), to precisely determine if these increases were related to oscillatory events or evoked power alone. However, as FOOOF has a higher frequency resolution requirement, we assessed for oscillations in 1-second windows (50ms to 1050ms) as opposed to the 500ms windows used previously. As such, we further restricted ourselves to the set of ROIs which exhibited significant low-frequency modulation in these same 1-second windows (lateral/medial orbitofrontal cortex, rostral anterior cingulate, isthmus of the cingulate, and precentral gyrus), in order to provide a fair comparison.

FOOOF was applied to power spectra from 3 to 50Hz, covering the theta and gamma ranges. We used a lower spectral peak width limit of 0.5 Hz and a upper limit of 12 Hz. Peaks were required to exceed 3 standard deviations of the aperiodic baseline in order to be considered as possible oscillatory peaks. To statistically compare the presence of oscillations in TMS versus sham trials, the average peak power was taken across all identified TMS events and sham events, for each electrode, and the results were subtracted from one another. Across all electrodes which fell within a region, the TMS-minus-sham difference was tested against zero using a 1-sample *t-* test, generating a new *t*-statistic reflecting the TMS-related increase (or decrease) in oscillatory power. The theta range was analyzed by considered peaks which fell within 3-8 Hz.

## Acknowledgments

We thank the neurosurgical patients who selflessly participated in this research. We thank Joel Bruss for assistance with imaging analyses. We also thank Christopher Kovach, Ariane Rhone, Haiming Chen, and Benjamin Pace for their assistance with image processing, coordinating, and conducting the experiments.

## Funding

NIMH R01MH132074 (CJK and ADB)

R21MH120441 and 5R01dC004290-20 (ADB)

R01NS114405 (ADB)

F30MH119763 (JBW)

Mark and Mary Stevens Interdisciplinary Graduate 660 Fellowship (JBW)

R01MH126639 (CJK)

R01MH129018 (CJK)

Burroughs Wellcome Fund Career Award for Medical Scientists (CJK)

5T32-MH019113 (NTT)

This work was conducted, in part, on an MRI instrument funded by 1S10OD025025-01

## Author contributions

EAS, JBW, HO, CJK, and ADB conceived of the study. EAS and JBW performed electrophysiology analyses. HO performed safety testing and analyses and participated in experimental testing with TMS. BDU and NTT participated in experimental testing. MH assisted with study design, safety, and subject recruitment. All authors contributed to the writing of the manuscript. ADB and CJK jointly supervised all aspects of the study. All authors have seen and approved the manuscript, and it has not been accepted or published elsewhere.

## Competing interests

CJK holds equity in Alto Neuroscience, Inc. EAS has previously received compensation for ad-hoc technical consulting for Nia Therapeutics, a company intended to develop and commercialize brain stimulation therapies. No other conflicts of interest, financial or otherwise, are declared by the authors.

## Data and materials availability

The de-identified raw data used in this study will be made available on the NIMH Data Archive upon publication. Preprocessing and analysis code has been made available in a public GitHub repository: <u>KellerLab-Stanford/Analysis-TMSiEEG</u> (github.com)

## Notes

### Summary of Updates

Added three new main text figures including (1) new analyses to differentiate theta oscillations from low-frequency evoked power, (2) subregional assessment of TMS-provoked effects, and (3) evaluation of cingulate responses to DLPFC stimulation.

