## Supplemental Figures and Tables for "TMS provokes target-dependent intracranial rhythms across human cortical and subcortical sites"

Ethan A. Solomon* *et al.*

**This PDF file includes:**

Figs. S1 to S3

Tables S1

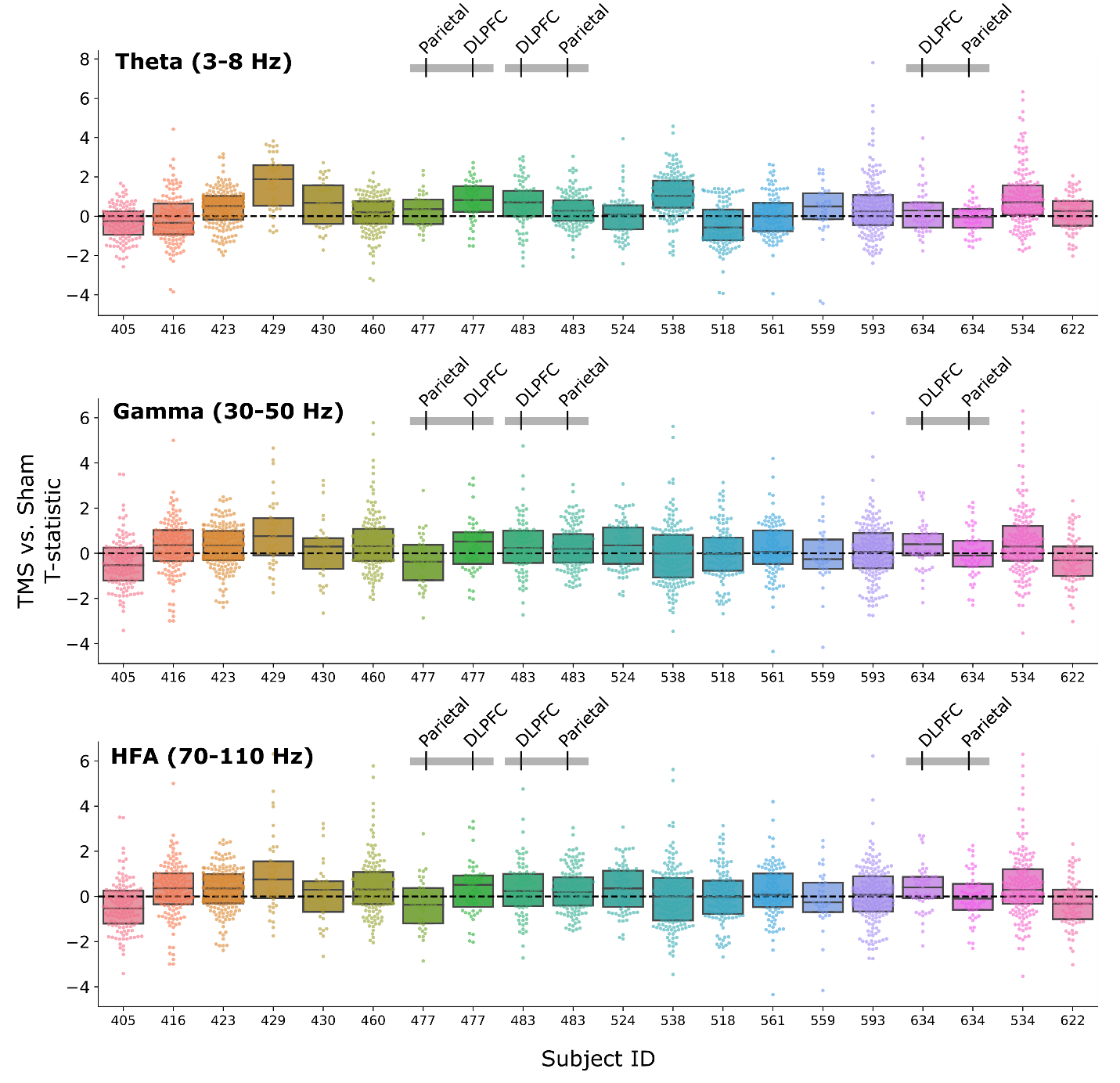

**Figure S1. TMS elicits brain-wide modulations of spectral power.** For each recording contact in the dataset, a *t*-statistic is computed which reflects the TMS-related change in spectral power relative to sham in the 50-500ms (theta, gamma) or 50-250ms (HFA) interval (see *Methods* and **Figure 1** for details). The distribution of *t*-statistics across all subjects and contacts is shown for each frequency band of interest (theta, gamma, and HFA). Three subjects (477, 483, 634) underwent stimulation at both parietal and DLPFC targets, indicated in the key above each plot. Boxes represent median and interquartile range.

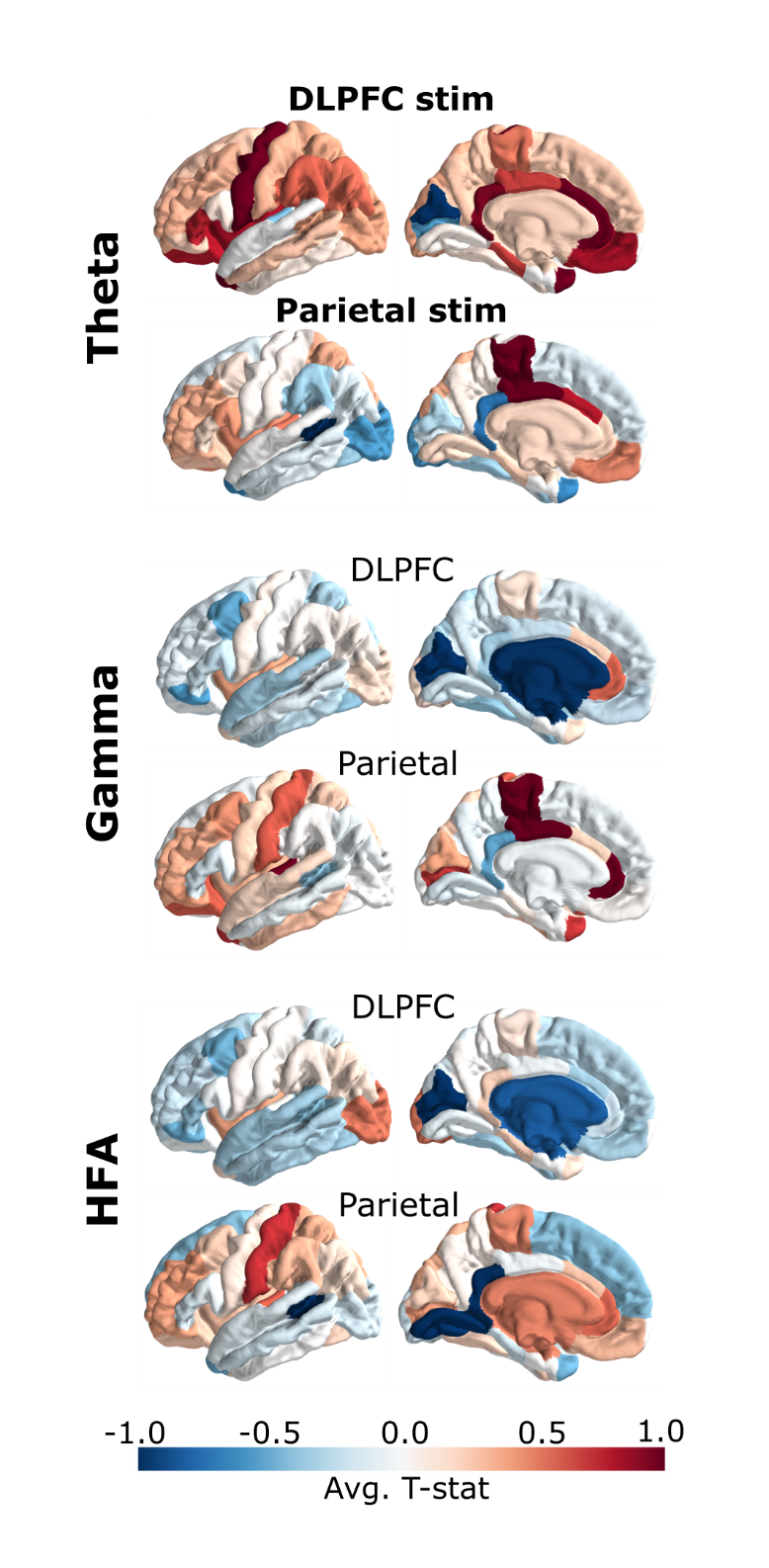

**Figure S2. Average TMS vs. Sham evoked power across DKT regions in complete dataset.** Exactly as **Figure** **3D**, but without a threshold for minimum number of subjects per ROI.

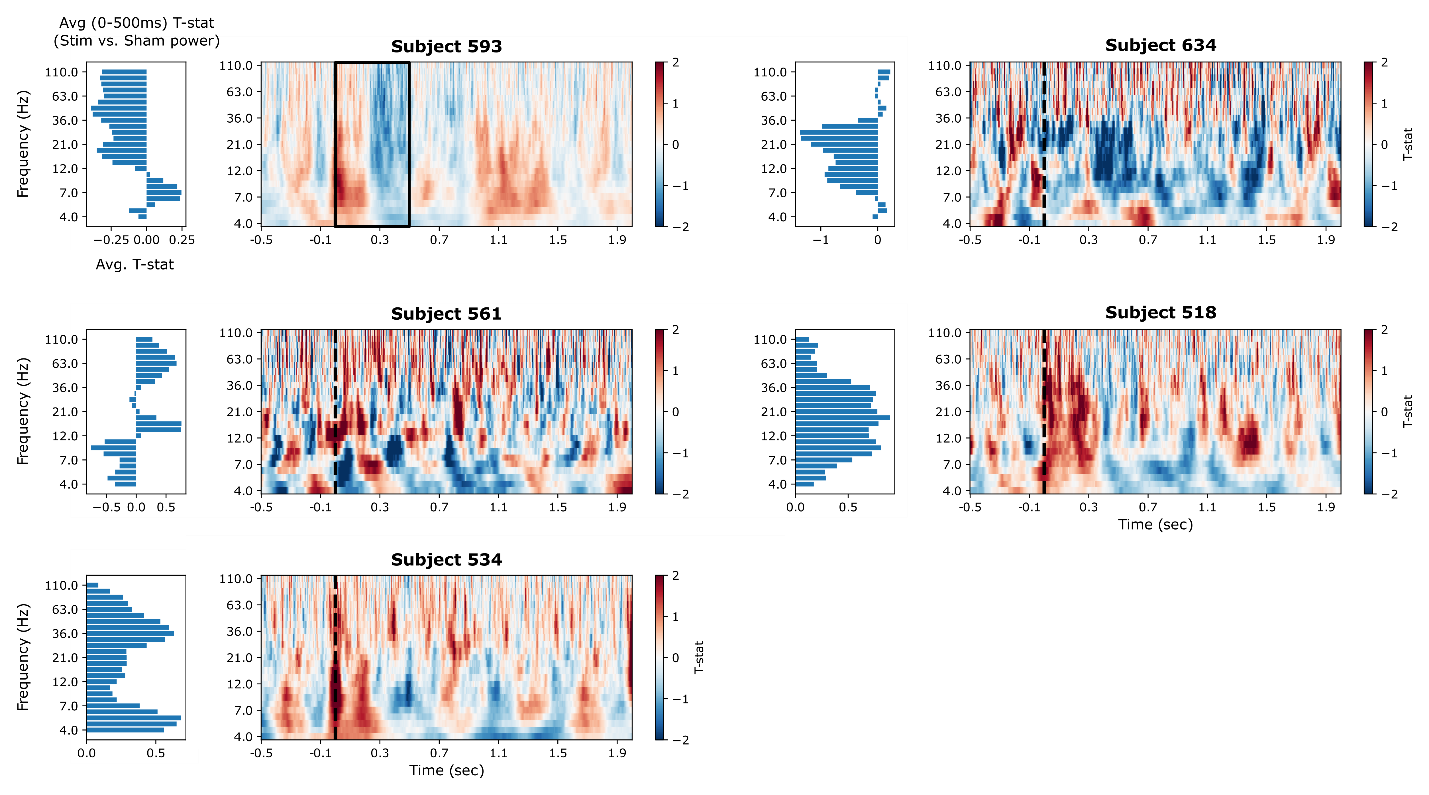

**Figure S3. Amygdala spectral power across individual subjects following DLPFC stimulation.** Time-frequency representations for each of the 5 subjects who had recording contacts in the amygdala and underwent TMS to the DLPFC. Time-frequency responses are constructed as described in **Figure 3A** and *Methods*. To the left of each time-frequency plot is the average power at each frequency within the 0-500ms post-stimulation interval. Subjects demonstrate heterogeneous effects with differing peak power for each subject; none show an entirely broadband response that would suggest contamination by stimulation artifact.

**Table S1. DKT regions included in broad ROIs.**

| **ROI** | **Constituent DKT labels** |
| --- | --- |
| Frontal | lateralorbitofrontal, parsorbitalis, precentral, caudalmiddlefrontal, parsopercularis, paracentral, rostralmiddlefrontal, medialorbitofrontal, superiorfrontal, parstriangularis, |
| Temporal | inferiortemporal, fusiform, temporalpole, superiortemporal, middletemporal, transversetemporal, bankssts |
| Parietal | supramarginal, inferiorparietal, superiorparietal, precuneus, cuneus |
| MTL | entorhinal, parahippocampal, Hippocampus |
| Limbic | entorhinal, insula, isthmuscingulate, posteriorcingulate, Amygdala, caudalanteriorcingulate, rostralanteriorcingulate, parahippocampal, Hippocampus |
